# Bayesian Shrinkage Estimation of High Dimensional Causal Mediation Effects in Omics Studies

**DOI:** 10.1101/467399

**Authors:** Yanyi Song, Xiang Zhou, Min Zhang, Wei Zhao, Yongmei Liu, Sharon L. R. Kardia, Ana V. Diez Roux, Belinda L. Needham, Jennifer A. Smith, Bhramar Mukherjee

**Affiliations:** Department of Biostatistics, University of Michigan, Ann Arbor, MI, U.S.A.; Department of Epidemiology, University of Michigan, Ann Arbor, MI, U.S.A.; Department of Epidemiology and Prevention, Wake Forest School of Medicine, Winston-Salem, NC, U.S.A.; Department of Epidemiology and Biostatistics, Drexel University, Philadelphia, PA, U.S.A.

## Abstract

Causal mediation analysis aims to examine the role of a mediator or a group of mediators that lie in the pathway between an exposure and an outcome. Recent biomedical studies often involve a large number of potential mediators based on high-throughput technologies. Most of the current analytic methods focus on settings with one or a moderate number of potential mediators. With the expanding growth of omics data, joint analysis of molecular-level genomics data with epidemiological data through mediation analysis is becoming more common. However, such joint analysis requires methods that can simultaneously accommodate high-dimensional mediators and that are currently lacking. To address this problem, we develop a Bayesian inference method using continuous shrinkage priors to extend previous causal mediation analysis techniques to a high-dimensional setting. Simulations demonstrate that our method improves the power of global mediation analysis compared to simpler alternatives and has decent performance to identify true non-null mediators. We also construct tests for natural indirect effects using a permutation procedure. The Bayesian method helps us to understand the structure of the composite null hypotheses. We applied our method to Multi-Ethnic Study of Atherosclerosis (MESA) and identified DNA methylation regions that may actively mediate the effect of socioeconomic status (SES) on cardiometabolic outcome.

## 1 Introduction

Causal mediation analysis has been of great interest across many disciplines [1, 2]. It investigates how an intermediate variable, referred to as mediator, explains the mechanism through which the exposure variable affects the outcome. Under certain regularity conditions, mediation analysis allows us to disentangle the exposure’s effect into two parts: effect that acts through the mediator of interest (indirect/mediation effect) and effect that is unexplained by the mediator (direct effect). The state-of-the-art causal mediation analysis [3, 4], which is built upon the counterfactual framework [5, 6], establishes rigorous assumptions regarding the exposure-outcome, exposure-mediator and mediator-outcome relationships to justify appropriate use of the classical formulas from Baron and Kenny in the linear regression setting [7, 8] and creates a framework for other general extensions. Many of the existing methods focus on univariate mediator analysis that analyzes one mediator at a time in the causal inference framework, and are applicable to both continuous [9] and binary outcomes [10], and also can account for exposure-mediator interactions [11]. These methods have been widely applied in areas of social, economic, epidemiological and genetic studies [4, 8], including recent extensions to multiple exposure variables that lead to more powerful single nucleotide polymorphism (SNP) set tests in presence of gene expression data [12]. Several studies have recently extended mediation analysis models to jointly account for multiple mediators. However, most of the literature considered settings with two or three mediators, where each mediator is ordered along a priori known mediation pathways and the path-specific effects are estimated [13, 14]. In the presence of multiple unordered mediators, one often has to rely on an *ad hoc* approach to fit a series of mediation models with one mediator and one exposure [15, 16]/outcome [17] at a time and then summarize the mediation effects across all the mediators. Such approach ignores correlation among mediators and the estimated mediation effect does not necessarily have an intuitive interpretation, particularly when the dimension of the potential mediators is truly large.

In this paper, building on the potential outcome framework for causal inference, we develop a Bayesian mediation analysis method in the presence of high-dimensional mediators. Bayesian methods for mediation have primarily been proposed in a principal stratification framework [18], in which the exposure effects on outcome are defined conditional on a single mediator. For estimating natural direct and indirect effects, recent work applied Bayesian non-parametric models, especially Dirichlet process mixture models [19, 20] in multiple mediators analysis. In contrast, here, we rely on Bayesian variable selection models to simultaneously analyze a relatively large number of mediators with potentially a small number being truly active. With sparsity inducing priors on mediator effects, we assume that only a small proportion of mediators may mediate the exposure effect on the outcome. This sparsity assumption allows us to extend previous univariate mediator analysis to a high-dimensional setting by casting the identification of active mediators as a variable selection problem and applying Bayesian methods with continuous shrinkage priors on the effects. Unlike previous methods developed for multiple mediators analysis, ours can simultaneously analyze much larger number of mediators without making any path-specific or causal ordering assumptions on mediators. Our method enables us to identify both the indirect effect of a specific mediator and the joint indirect effects of all the mediators, and propagates uncertainty in inference in a principled way.

While our method is generally applicable to many settings, we examine the performance of our method in the setting of genomics studies. Due to fast advances in high-throughput biological technologies, genomics studies can nowadays measure a large number of molecular-level traits such as gene expression and DNA methylation (DNAm) levels. Recent studies have proposed these molecular traits may act as a mechanism through which various aspects of socioeconomic status (SES) and neighborhood disadvantages affect physical health. For example, childhood SES, adult SES, social mobility, and neighborhood crime rates have recently been shown to influence DNAm in several genes related to stress and inflammation [21, 22]. DNAm of inflammatory markers have also been associated with the status of cardiovascular disease (CVD) [23] and type 2 diabetes (T2D) [24]. Here, we show through simulations and data analysis that our high-dimensional mediation analysis framework can increase power of a joint analysis and facilitate the identification of individual mediators.

## 2 Notation, Definitions and Assumptions

In this paper, we focus on causal mediation analysis for the setting where there is a single exposure of interest but a high-dimensional set of candidate mediators that may mediate the effect of exposure on an outcome. Suppose our analysis is based on a study of *n* subjects and for subject *i*, *i* = 1, …, *n*, we collect data on exposure *A_i_*, *p* candidate mediators ***M_i_***. = 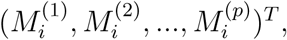 outcome *Y_i_*, and *q* covariates 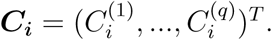 In particular, we focus on the case where *Y_i_* and ***M_i_*** are all quantitative variables.

We adopt the counterfactual (or potential outcomes) framework to formally define mediators and their causal effects. Let the vector 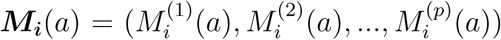 denote the *i*th subject’s potential (or counterfactual) value of the *p* mediators if, possibly contrary to fact, he/she received exposure *a*. Let *Y_i_*(*a*, ***m***) denote the ith subject’s potential outcome if the subject’s exposure were set to *a* and mediators were set to ***m***. These potential (or counterfactual) variables are hypothetical variables and may not be observed in real data. To connect potential variables to observed data, we make the Stable Unit Treatment Value Assumption (SUTVA) [25, 26], which is a commonly made assumption for performing causal inference. Specifically, the SUTVA assumes there is no interference between subjects and the consistency assumption, which states that the observed variables are the same as the corresponding potential variables when their determinants are set to the observed, i.e., ***M_i_*** = ***M_i_***(*a* = *A_i_*), and *Y_i_* = *Y_i_*(*a* = *A_i_*, ***m*** = ***M_i_***). For simplicity in notation, we define *Y_i_*(*a*) = *Y_i_*(*a*, ***M_i_***(*a*)), i.e., the potential outcome when exposure were set to *a* and mediators were set to the value that would have been observed had the exposure were set to *a*. Although potential or counterfactual variables are useful concepts in order to formally define causal effects, they are hypothetical and actually most of them are not observed in real data. For example, if *A_i_* ≠ *a*, then *Y_i_*(*a*) or *M_i_*(*a*) are not observed. Also *Y_i_*(*a*) and *Y_i_*(*a*^⋆^) are never simultaneously observed for a subject.

We may decompose the effect of an exposure into its direct effect and effect mediated through mediators. The controlled direct effect (CDE) of the exposure on the outcome is defined as *Y_i_*(*a*, ***m***) − *Y_i_*(*a*^⋆^, ***m***), which is the effect of changing exposure from level *a*^⋆^ (the reference level) to *a* while hypothetically controlling mediators at level ***m***. The natural direct effect (NDE) is defined as *Y_i_*(*a*, ***M_i_***(*a*^⋆^)) − *Y_i_*(*a*^⋆^, ***M_i_***(*a*^⋆^)), which is the CDE when mediators are controlled at the level that would have naturally been had the exposure been *a*^⋆^. The natural indirect effect (NIE) is defined by *Y_i_*(*a*, ***M_i_***(*a*)) − *Y_i_*(*a*, ***M_i_***(*a*^⋆^)), capturing the part of exposure effect mediated through mediators, i.e., the change in potential outcomes when mediators change from ***M_i_***(*a*^⋆^) to ***M_i_***(*a*) while fixing exposure at *a*. The total effect (TE), *Y_i_*(*a*) − *Y_i_*(*a*^⋆^), can then be decomposed into natural direct effect and natural indirect effect, written as *Y_i_*(*a*) − *Y_i_*(*a*^⋆^) = *Y_i_*(*a*, ***M_i_***(*a*)) − *Y_i_*(*a*^⋆^, ***M_i_***(*a*^⋆^)) = *Y_i_*(*a*, ***M_i_***(*a*)) − *Y_i_*(*a*, ***M_i_***(*a*^⋆^)) + *Y_i_*(*a*, ***M_i_***(*a*^⋆^)) − *Y_i_*(*a*^⋆^, ***M_i_***(*a*^⋆^)) =NIE+NDE.

Causal effects are formally defined in terms of potential variables which are not necessarily observed, but the identification of causal effects must be based on observed data. Therefore, similar to missing data problems, further assumptions regarding the confounders are required for the identification of causal effects in mediation analysis [15]. We will use *A* ╨ *B*∣*C* to denote that *A* is independent of *B* conditional on *C*. For estimating the average CDE, two assumptions on confounding are needed: (1) *Y_i_*(*a*, ***m***)╨*A_i_*∣***C_i_***, namely, there is no unmeasured confounding for the exposure effect on the outcome; (2) *Y_i_*(*a*, ***m***)╨***M_i_***∣{***C_i_***, *A_i_*}, namely, there is no unmeasured confounding for any of mediator-outcome relationship after controlling for the exposure. The two assumptions are illustrated in the left panel of Figure 1, and controlling for exposure-outcome and mediator-outcome confounding corresponds to controlling for *C*_1_, *C*_2_ in the figure. In practice, both sets of covariates *C_1_* and *C*_2_ need not to be distinguished from one another and can simply be included in the overall set of *C* that we adjust for. The identification of the average NDE and NIE requires assumption (1) and (2), along with two additional assumptions: (3) ***M_i_***(*a*)╨*A_i_*∣***C_i_***, namely, there is no unmeasured confounding for the exposure effect on all the mediators; (4) *Y_i_*(*a*, ***m***)╨***M_i_***(*a*^⋆^)∣***C_i_***, which can be interpreted as there is no downstream effect of the exposure that confounds the mediator-outcome relationship for any of the mediators. Graphically, assumption (4) implies that there should be no arrow going from exposure *A* to mediator-outcome confounder *C*_2_ in Figure 1(a). It is thus violated in Figure 1(b) since the mediator-outcome confounder *L* is itself affected by the exposure. The four assumptions are required to hold with respect to the whole set of mediators ***M_i_***(*a*) = 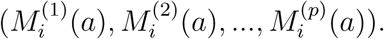 Finally, as in all mediation analysis, in order for associations to represent causal effects, the temporal ordering assumption also needs to be satisfied, i.e., the exposure precedes the mediators and the mediators precede the outcome.

**Figure 1:**
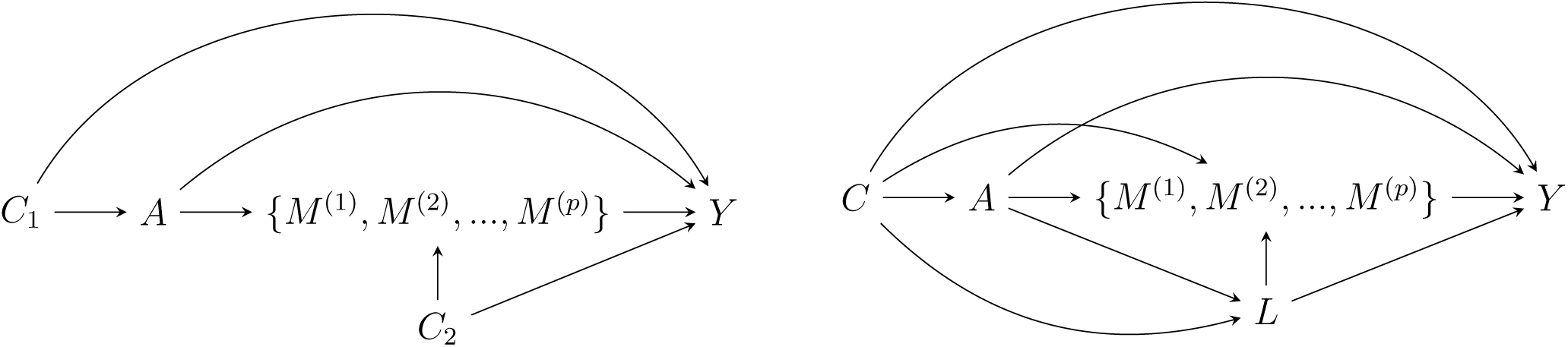
Left (a): High-dimensional mediators ((*M*^(1)^,*M*^(2)^, …,*M*^(*p*)^)) between exposure (*A*) and outcome (*Y*) with exposure-outcome confounders *C*_1_ and mediator-outcome confounders *C*_2_; Right (b): An example of mediator-outcome confounder *L* that is affected by the exposure *A*.

Now we show that if the above assumptions hold, then the average natural direct and indirect effects can be identified from the observed data. We first notice that *E*[*Y_i_*(*a*, ***M_i_***(*a*^⋆^)∣***C_i_***] can be expressed as below (see Supplementary Materials for details),

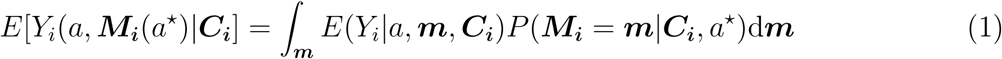

If we replace *a* with *a*^⋆^ in *E*[*Y_i_*(*a*, ***M_i_***(*a*^⋆^)∣***C_i_***], then we get *E*[*Y_i_*(*a*^⋆^, ***M_i_***(*a*^⋆^)∣***C_i_***] = *∫_m_ E*(*Y_i_*∣*a*^⋆^, ***m***, ***C_i_***)* *P*(***M_i_*** = ***m***∣***C_i_***,*a*^⋆^)d***m***. Therefore, we can express the average natural direct effect conditional on *C* as,

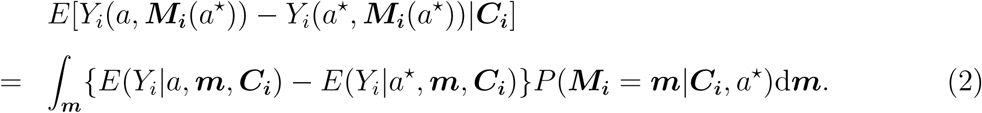

If we replace *a*^⋆^ with *a* in *E*[*Y_i_*(*a*, ***M_i_***(*a*^⋆^)∣***C_i_***], then we get *E*[*Y_i_*(*a*, ***M_i_***(*a*)∣***C_i_***] = *∫_m_ E*(*Y_i_*∣*a*, *m*, ***C_i_***)* *P*(***M_i_*** = ***m***∣***C_i_***, *a*)d***m***, and thus the average indirect effect conditional on *C* is given by,

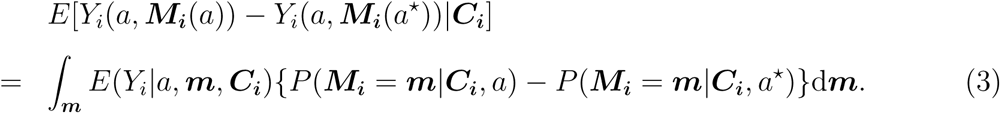

Finally, one can get the average NDE and NIE by taking expectation over *C* of the two conditional effects defined in (2) and (3). Importantly, Equations (2), (3) show that, under the assumptions we made, the average NDE and the average NIE can be identified by modeling *Y_i_*∣*A_i_*, ***M_i_***, ***C_i_*** and ***M_i_***∣*A_i_*, ***C_i_*** using observed data.

## 3 Models and Estimands

As discussed in Section 2, effects of mediators (average NDE and NIE) defined in terms of potential outcomes can be deduced from two conditional models for *Y_i_*∣*A_i_*, ***M_i_***, ***C_i_*** and ***M_i_***∣*A_i_*, ***C_i_*** using observed data. Therefore, we propose two regression models for the two conditional relationships and subsequently deduce the causal effects of mediators. For modeling *Y_i_*∣*A_i_*, ***M_i_***, ***C_i_***, we assume for subject *i*(*i* = 1, …, *n*), a continuous outcome of interest *Y_i_* is associated with exposure *A_i_*, *p* potential mediators ***M_i_*** = 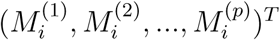 that may be on the pathway from *A_i_* to *Y_i_*, and *q* covariates ***C_i_*** with the first element being the scalar 1 for the intercept:

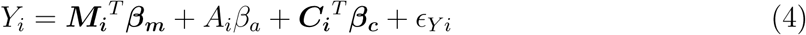

where ***β_m_*** = (*β*_*m*1_, …,*β_mp_*)^*T*^, ***β_c_*** = (*β*_*c*1_, …,*β_cq_*)^*T*^, 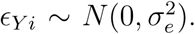 Here we assume there is no interaction between *A_i_* and ***M_i_***. Next for modeling ***M_i_***∣*A_i_*, ***C_i_*** we consider a multivariate regression model that jointly analyzes the *p* potential mediators:

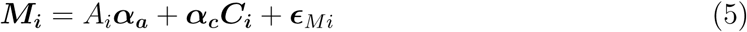

where ***α_a_*** = (*α*_*a*1_, …, *α_ap_*), ***α_c_*** (***α*_*c*1_**^*T*^, …, ***α_cp_***^*T*^)^*T*^, ***α***_*c*1_, … ***α_cp_*** are *q*-by-1 vectors, ***ϵ***_*Mi*_ ~ *MVN*(**0**, Σ), Σ captures the correlation among the mediators. *ϵ_Yi_* and ***ϵ_Mi_*** are assumed independent of *A_i_*, ***C_i_*** and each other.

With assumptions made in Section 2 and under the regression models specified for the outcome *E*(*Y_i_*∣*A_i_*, ***M_i_***, ***C_i_***) and for the mediators *P*(***M_i_*** ∣*A_i_*, ***C_i_***), we can analytically calculate the right-hand side of Equations (2), (3). We show in Supplementary Materials that the average NDE, NIE and TE can then be computed as below, and in the rest of the paper, we refer to NDE as direct effect and NIE as indirect/mediation effect.

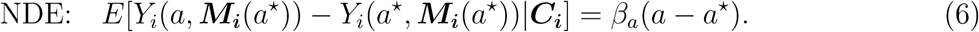

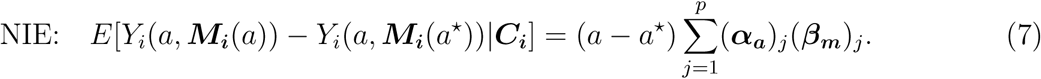

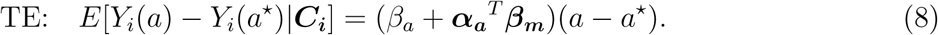

According to Equation (7), to select active mediators among the potential high-dimensional set of mediators is equivalent to identify the ones with marginsl indirect effect (***α_a_***)_*j*_ (***β_m_***)_*j*_ being non-zero. On the other hand, any inactive (un-selected) mediator will naturally fall into one of the following three categories: (***β_m_***)_*j*_ is non-zero while (***α_a_***)_*j*_ is zero; (***α_a_***)_*j*_ is nonzero while (***β_m_***)_*j*_ is zero; both are zero. Such a refined partition for the high-dimensional set of mediators provides useful and insightful interpretations on the way in which a mediator links or does not link exposure to outcome, and furthermore facilitates understanding the composite of non-mediating cases.

Regarding a global measure of the indirect effects, we note that the quantity in Equation (7), summation of each mediator’s marginal mediation effect, is a good summary of the global mediation effects when the marginal mediation effect for each mediator is of the same direction. However, when marginal mediation effects have opposite directions, their effects may cancel out and result in a small or zero indirect effect. Considering this, we propose to use the *L*_2_ norm of the vector of marginal mediation effects [17] as a global measure of mediation effects, i.e.,

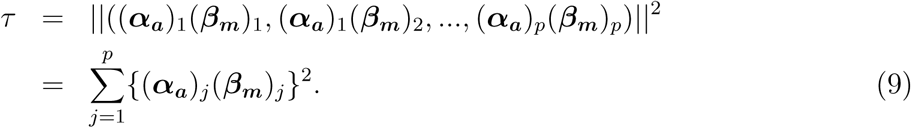

Such quadratic quantity, which involves a sum of squared terms, has been widely studied for testing associations between rare genetic variants and complex human traits, and shown to have reasonable power across a wide range of alternatives with correlated predictor set [27].

## 4 Bayesian Method for Estimation

### 4.1 Prior Specification

In order to conduct high-dimensional mediation analysis, we need to make certain model assumptions on the effect sizes. In genome-wide association studies, Bayesian sparse regression models, such as Bayesian variable selection regression models (BVSR), have been proven to yield better power in detecting relevant covariates [28]. For high-dimensional mediation analysis, we also make the reasonable sparsity assumption, which implies that only a small proportion of mediators mediate the exposure effects on the outcome. In practice, the exposure effects on the mediators and the mediator effects on the outcome may be small but not exactly zero. Linear mixed models (LMM), on the other hand, assume that every mediator transmits certain effects from exposure to outcome, with the effect sizes normally distributed. Therefore, in this paper, we propose Baysian Sparse Linear Mixed Model (BSLMM), a hybrid between LMM and BVSR [29] that imposes continuous shrinkage on the effects, for high-dimensional mediation analysis. The BSLMM is capable of learning the underlying mediation architecture from the data, producing good performances across a wide range of scenarios. Specifically, we assume a mixture of two normal components a priori for the *j*th mediator, *j* = 1, 2, …*p*,

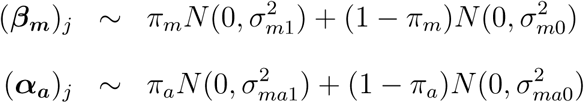

where 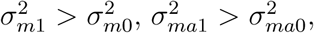 and *π_m_*, *π_a_* denote the proportion of coefficients that belong to the normal distribution with a larger variance.

For the other coefficients, we assume,

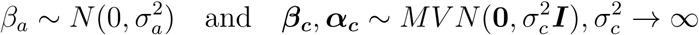

Here we use a limiting normal prior for ***β_c_***, ***α_c_*** with its variance going to infinity, since we often have sufficient information from the data to overwhelm any prior assumptions. For the convenience of modeling, we set the correlation structure among mediators **Σ** as 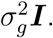

For the hyper-parameters of variances in the model, we use the standard conjugate priors,

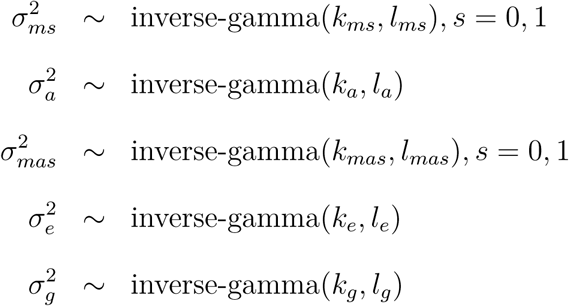

We set *k*_*m*0_ = *k*_*m*1_ = *k_a_* = *k*_*ma*0_ = *k*_*ma*1_ = *k_e_* = *k_g_* = 2.0, and *l*_*m*0_ = *l*_*ma*0_ = 10^−4^, *l_a_* = *l*_*m*1_ = *l*_*ma*1_ = *l_e_* = *l_g_* = 1.0. Following [29], we place a uniform prior on log(*π_m_*),log(*π_a_*),:

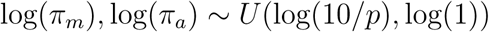

where *p* is the number of mediators. The priors were chosen so that *π_m_* and *π_a_* range from 10/*p* to 1, and the lower and upper bounds correspond to an expectation of 10 and *p* covariates in each model. A uniform prior on log(*π_m_*) and log(*π_a_*) reflects the fact that the uncertainty in *π_m_*,*π_a_* spans orders of magnitude.

### 4.2 Posterior Sampling Algorithm

We develop a Markov chain Monte Carlo (MCMC) sampling algorithm to obtain the posterior samples from our Bayesian method. To facilitate MCMC, we introduce indicator variables *r_m_*, *r_a_* ∈ {0,1}^*p*^ to indicate which normal component (*β_m_*)*_j_* and (*α_a_*)_*j*_ come from, and for the *j*th mediator, 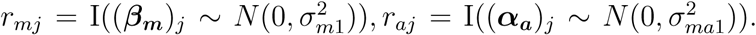 Let = 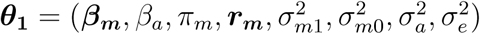 denote all the unknown parameters in model (4), and 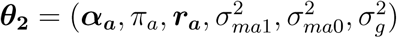 for model (5). The joint log posterior distribution is,

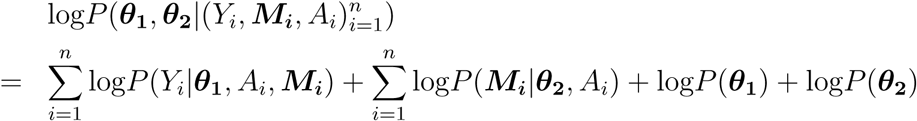

We use a Hastings-within-Gibbs algorithm to obtain posterior samples, and full details of the sampling algorithm appear in Supplementary Materials.

For the *j*th mediator, we can estimate the posterior probability of both (*β_m_*)*_j_* and (*α_a_*)_*j*_ being in the normal components with larger variances as the posterior inclusion probability (PIP), defined as *P*(*r_mj_* = 1,*r_aj_* = 1|Data) in our model. The PIP estimated in this way measures the association strength between exposure and mediators in model (5) and between mediators and outcome in model (4). Therefore, we select mediators with the highest PIP as the potentially active mediators.

### 4.3 Mediator Categorization

Under the above Bayesian mediation framework, active mediators are the ones whose (*β_m_*)_*j*_ and (*α_a_*)_*j*_ both come from larger normal components. The three categories for the inactive mediators are: (*β_m_*)*_j_* from larger normal component while (*α_a_*)*_j_* from smaller normal component; (*α_a_*)*_j_* from larger normal component while (*β_m_*)*_j_* from smaller normal component; both from smaller components. In addition to identifying true mediators, our method automatically classifies all the mediators into four groups based on their relationship with exposure and outcome. In practice, we have the indicator variables *r_mj_* and *r_aj_* to denote which component the coefficients (*β_m_*)*_j_*, (*α_a_*)_*j*_ belong to and can easily obtain the posterior probabilities for each group. The four groups are illustrated in Table 1,

**Table 1:** Mediators are categorized into four groups based on their relationships with exposure and outcome; Group 1: Both (*β_m_*)*_j_* and (*α_a_*)*_j_* come from larger normal components; Group 2: (*α_a_*)*_j_* from larger normal component while (*β_m_*)*_j_* from smaller normal component; Group 3: (*β_m_*)*_j_* from larger normal component while (*α_a_*)*_j_* from smaller normal component; Group 4: Both (*β_m_*)_*j*_ and (*α_a_*)*_j_* come from smaller normal components.

| | $(\beta_m)_j$ | Large component | Small component |
| --- | --- | --- | --- |
| $(\alpha_a)_j$ | | | |
| Large component | | $r_{mj} * r_{aj} = 1$ (Group 1) | $r_{mj} = 0, r_{aj} = 1$ (Group 2) |
| Small component | | $r_{mj} = 1, r_{aj} = 0$ (Group 3) | $r_{mj} = r_{aj} = 0$ (Group 4) |

### 4.4 Global Test of Mediation Effects

Typically the majority of mediators are not actively mediating the exposure effect on the outcome, so it is natural to focus on the global null hypothesis and test for *H*_0_ : *τ* = 0,

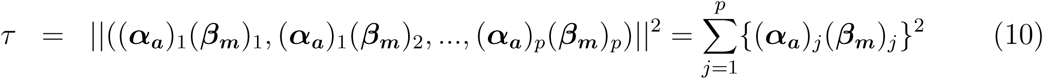

One estimate for *τ* from our method is the sampling mean calculated from the posterior samples, 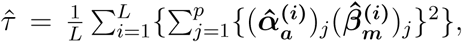 where 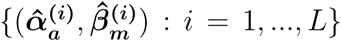 are the *L* samples generated from posterior distributions.

It is difficult to analytically derive the composite null distribution of *τ* because the null distribution depends on the proportion of mediators belonging to each of the four categories. Instead, we resort to the permutation method. The global null indicates that none of the mediators are active, and to construct the three null components for each mediator, the following permutation procedures are brought up: (a) permute 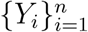 in Equation (4) to dissolve the relationship between outcome and mediators and obtain the null estimates of 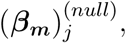 (b) permute 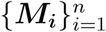 in Equation (5) to dissolve the relationship between mediators and exposure and obtain the null estimates of 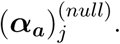 We then propose the following three quantities: (i) 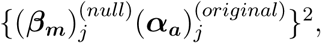 (ii) 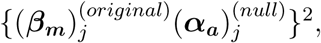 (iii) 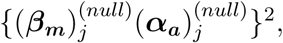 where 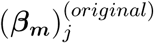 and 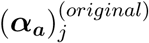 are estimated from the original data. For the jth mediator, (i), (ii) and (iii) correspond to the three components of the null distribution: *r_aj_* = 1,*r_mj_* = 0 (Group 2), *r_mj_* = 1,*r_aj_* = 0 (Group 3), *r_mj_* = 0,*r_aj_* = 0 (Group 4), respectively. Each quantity is then weighted by the posterior probability for that group estimated from original data, and we finally sum the results across all the mediators to obtain draws from the empirical null distribution of *τ*.

## 5 Simulations

We evaluate the performance of the proposed Bayesian mediation method and compare it with two other existing mediation methods in our simulations. The two existing methods include single mediation analysis and multivariate mediation analysis. Single mediation analysis tests one mediator at a time for its mediation effect. We use the R package **mediation** to run single mediation analysis with the nonparametric bootstrap option for standard error estimation. Multivariate mediation analysis [15], on the other hand, jointly analyzes all the mediators in both model (4) and (5) and tests the product term (***β_m_***)_*j*_ (***α_a_***)_*j*_ for each *j* at a time while controlling for all other variables. This method can only be fit when a multivariate ordinary least squares regression model can be fit for the outcome model (4). We implement the multivariate mediation analysis and compute the standard error based on first and second order Taylor series approximation of the product [30]: 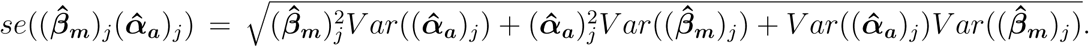 Afterwards, we obtain a *z*-statistics for the *j*th mediator by dividing 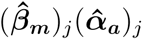 with its standard error and compute the corresponding *p*-value based on asymptotic normality. We use *p*-values for the two existing frequentist methods (univariate and multivariate) and PIP for our Bayesian method as measures of the evidence for mediation. We compare the power to identify active mediators based on either 5% or 10% false discovery rate (FDR).

We consider various simulation settings with *n* = 1,000 samples and *p* mediators (*p* = 100 or 2,000). Since the multivariate mediation analysis can only be applied to settings where the number of mediators is smaller than the number of observations (i.e. *p* < *n*), we first examine the settings where *p* = 100 in order to include the multivariate mediation analysis for comparison. We will later consider the high-dimensional setting where *p* = 2,000. For each simulation setting, we first simulate a set of continuous exposure variables {*A_i_*, *i* = 1, …, 1000} independently from a standard normal distribution. We then generate a *p*-vector of mediators for the *i*th individual from ***M_i_*** = *A_i_**α_a_*** + ***ϵ**_Mi_*. Here, each element of ***α_a_***, (***α_a_***)_*j*_ (*j* = 1, …, *p*), is simulated from a point-normal prior: *π_a_N*(0,1)+(1–*π_a_*)*δ*_0_, where *δ*_0_ is a point mass at zero. The residual errors ***ϵ**_M i_* are simulated from a multivariate normal distribution with mean zero and a covariance ∑. ∑ accounts for the correlation among mediators commonly seen in real data, and we use the sample covariance estimated from the Multi-Ethnic Study of Atherosclerosis (MESA) data to serve as ∑. Because our Bayesian mediation model does not explicitly account for the correlation structure of mediators in the model between mediators and exposure, the simulations with correlated mediators allow us to examine the robustness of our modeling assumption regarding independence. After simulating *A_i_**α_a_*** and ***ϵ**_M i_*, we scale these two terms further so that the first term explains a fixed proportion of variance: *PVE_A_* = *V ar*(*A_i_**α_a_***)/*V ar*(***M_i_***), where *V ar* denotes the sample variance.

Given the exposure and mediators, we then generate the outcome *Y_i_* from the linear model: 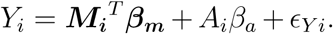 Here, each element of ***β_m_***, (***β_m_***)_*j*_ (*j* = 1, …, *p*), is simulated from *π_m_N*(0, 1) + (1 – *π_m_*)*δ*_0_, where *δ*_0_ is a point mass at zero, and *β_a_* from a standard normal distribution. The residual error *ϵ_Y i_* is simulated independently from a standard normal distribution. We assume that only 10% of the mediators are truly mediating the exposure effects on the outcome (i.e. active mediators), whose (***β_m_***)_*j*_ and (***α_a_***)_*j*_ are both sampled from the large variance normal distribution. After simulating 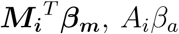 and *ϵ_Yi_*, we scale these three terms further to achieve two desirable *P V E*s: *P V E_IE_* = *V ar* 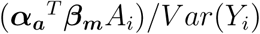 and *P V E_DE_* = *V ar*(*A_i_β_a_*)/*V ar*(*Y_i_*).

To explore a variety of simulation scenarios, we first examine a baseline scenario where we set *P V E_A_* = 0.5, *P V E_IE_* = 0.4, *P V E_DE_* = 0.1, *π_a_* = 0.3, *π_m_* = 0.2. We then vary each of the four parameters (*P V E_A_*, *P V E_IE_*, *π_a_*, *π_m_*) one at a time in different scenarios to investigate their individual influences on the results. We perform 200 replicates for each simulation scenario to do the power comparison.

We first examine the settings for *p* = 100 and display the comparative results in Figure 2. The results show that our Bayesian multivariate mediation method outperforms both the univariate and multivariate mediation analysis methods in all scenarios. For example, in the baseline setting, at 10% FDR, Bayesian mediation method achieves a power of 0.725, while the univariate and multivariate methods achieve a power of 0.527 and 0.676, respectively. The power of the three approaches increases with increasing *PVE_IE_*, which increases the effect sizes of *β_m_*. In addition, the power of various approaches reduces with increased *π_a_* or *π_m_*, which reduces the effect sizes of either *α_a_* or *β_m_*, respectively. As one would expect, the advantage of our Bayesian method over the other two methods is more apparent in sparse settings with smaller values of *π_a_* and *π_m_*. In terms of *PVE_A_*, which determines the effect size of *α_a_*, we found that the power of different methods first increases slightly when *PVE_A_* changes from 0.3 to 0.5 and then decreases slightly as *PVE_A_* changes further to 0.8. The later decrease in power in the setting of *PVE_A_* = 0.8 is presumably due to the increased correlation between the exposure and mediators, which makes it difficult for all the three methods to distinguish between direct effects and indirect effects in model (4). Between the two competing methods, the multivariate mediation analysis method yields better power than the single mediation analysis method in all scenarios, as the multivariate mediation analysis properly controls for the correlation among mediators.

**Figure 2:**
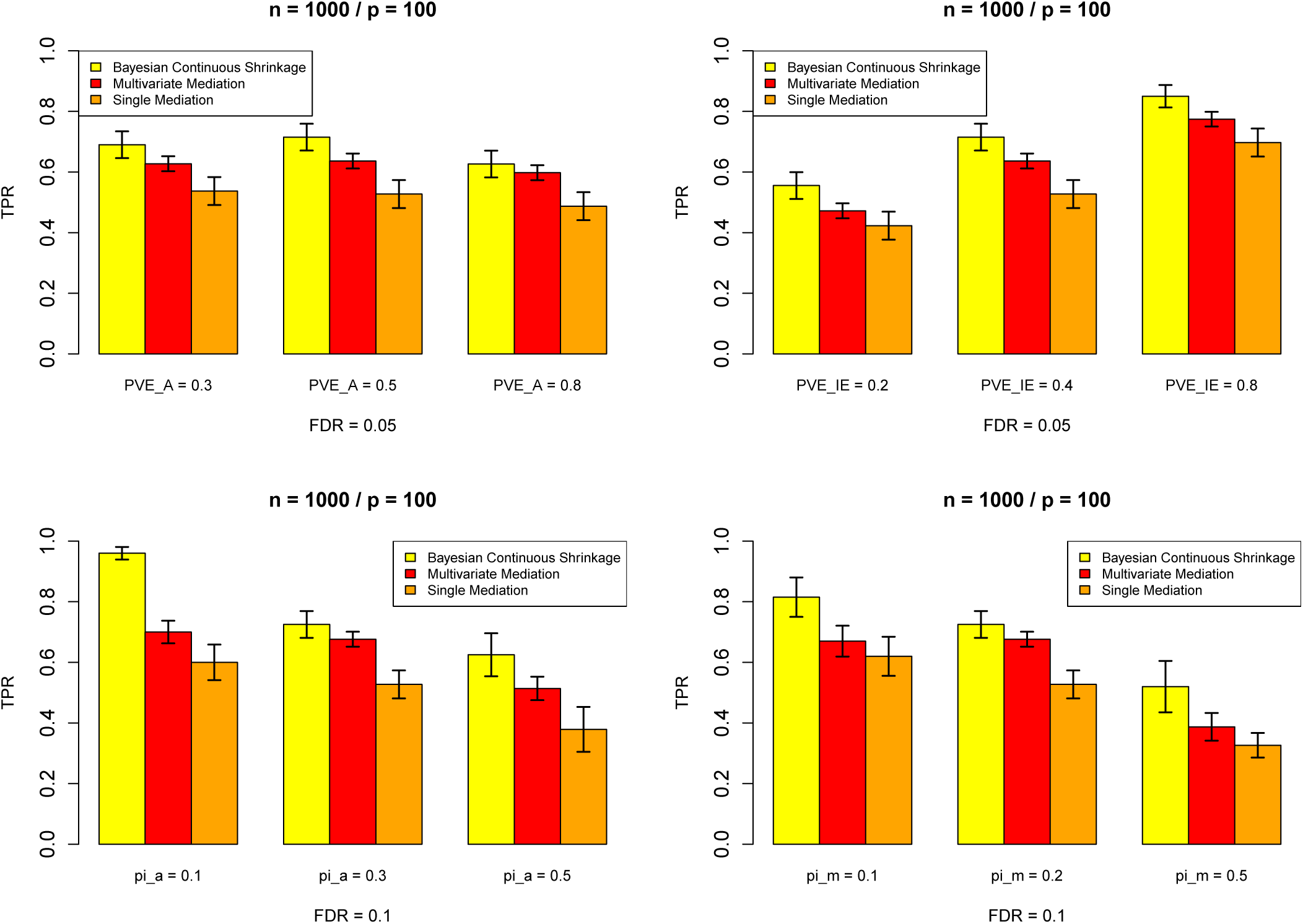
Power comparison among our Bayesian mediation method (yellow), multivariate mediation method (red) and single mediation method (orange) when the number of mediators is 100 and sample size 1,000. The x-axis marks the one parameter we change at a time from the baseline setting. The average TPR at FDR = 0.05/0.1 and its error bar are calculated across 200 replicates.

Next, we examine the settings for *p* = 2, 000. Now we select 1% of the mediators to be active and set *π_m_* = 2%, *π_a_* = 3% as the baseline setting with all other configurations being same as in the baseline setting of *p* = 100. Since the multivariate mediation analysis is unfeasible when *p* > *n*, we compare our method with single mediation analysis alone. We use a threshold of 1% false positive rate (FPR) instead of false discovery rate due to low power in the *p* = 2, 000 settings. The comparisons are shown in Figure 3. The Bayesian mediation method yields more power than the single mediator analysis in all scenarios. For example, in the baseline setting, at 1% FPR, Bayesian mediation method achieves a power of 0.470, while the univariate method has a power of 0.357. The power of the two approaches again increases with increasing *PVE_E_* and decreases with increasing *π_a_* or *π_m_*. The power of the Bayesian method decreases with increasing *PVE_A_*, while the power of the univariate analysis method is relatively stable. In addition, our Bayesian method with continuous shrinkage is more powerful than the univariate method especially when the model is most sparse.

**Figure 3:**
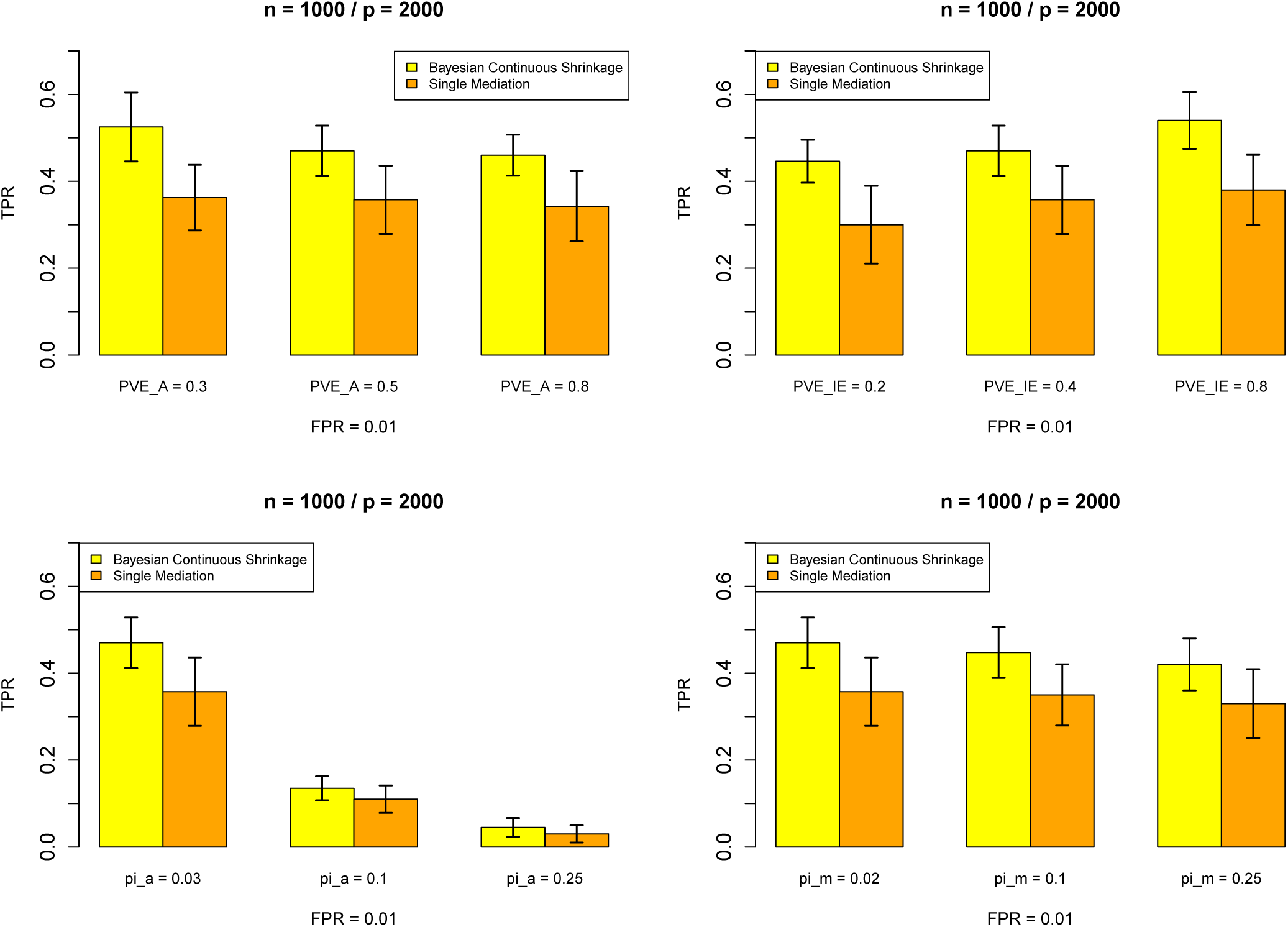
Power comparison between our Bayesian mediation method (yellow) and single mediation method (orange) when the number of mediators is 2,000 and sample size 1,000. The x-axis marks the one parameter we change at a time from the baseline setting. The average TPR at FPR = 0.01 and its error bar are calculated across 200 replicates.

Finally, we examine the ability of our method to detect the overall mediation effect and estimate the proportion of mediators in the four different categories. The four categories are characterized by the effect of the mediator on the outcome and the effect of the exposure on the mediator as shown in Table 1. We use *π*_*g*1_, *π*_*g*2_, *π*_*g*3_, *π*_*g*4_ to represent the proportion of mediators in Group 1, Group 2, Group 3 and Group 4, respectively. We examine eight different simulation scenarios based on different combinations of *π*_*g*1_, *π*_*g*2_, *π*_*g*3_ and *π*_*g*4_, which include four null scenarios with *π*_*g*1_ = 0 and four alternative scenarios with *π*_*g*1_ ≠ 0. In these simulations, we set *PVE*_s_ to be the same as in the baseline setting (*PVE_A_* = 0.5, *PVE_IE_* = 0.4, *PVE_DE_* = 0.1; except when *π*_*g*4_ = 1 where *PVE_A_* and *PVEI_E_* are zero).

To detect the overall mediation effect, we obtain the posterior mean of *τ*. For each simulated data, we apply the permutation procedure described in Section 4.4 with 100 permutations and compare the estimated 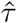 with the empirical null distribution. We find that the global test controls well the Type I error under four different null scenarios (Table 2). In addition, the global test yields reasonable power under the four different alternative scenarios (Table 3). In most scenarios, the power is close to or above 0.8. Finally, we estimate the proportion of mediators in each of the four different categories using posterior samples and find that our method provides decent estimates for *π*_*g*1_, *π*_*g*2_, *π*_*g*3_ and *π*_*g*4_ across different scenarios (Tables 2 and 3). Note that our estimates for *π*_*g*1_, *π*_*g*2_, *π*_*g*3_ are slightly conservative due to the fact that our model does not have full power to detect all the mediators.

**Table 2:** Empirical size of the proposed global test at level of 0.01 based on 1,000 simulations when *p* = 100/2, 000. We denote *π*_*g*1_, *π*_*g*2_, *π*_*g*3_ and *π*_*g*4_ to represent the proportion of mediators in Group 1, Group 2, Group 3 and Group 4 as defined in Table 1, and 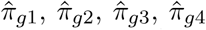 are the estimated proportions from our Bayesian method. The true *PVE_IE_* = 0.4, and 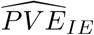 is the estimated value.

| $p$ | Composition | $\pi_{g1}$ | $\pi_{g2}$ | $\pi_{g3}$ | $\pi_{g4}$ | Size | $\hat{\pi}_{g1}$ | $\hat{\pi}_{g2}$ | $\hat{\pi}_{g3}$ | $\hat{\pi}_{g4}$ | $\widehat{PVE}_{IE}$ |
| --- | --- | --- | --- | --- | --- | --- | --- | --- | --- | --- | --- |
| 100 | A | 0 | 0.2 | 0.1 | 0.7 | 0.005 | 0.003 | 0.195 | 0.074 | 0.728 | 0.397 |
|  | B | 0 | 0.1 | 0.2 | 0.7 | 0.003 | 0.003 | 0.097 | 0.115 | 0.786 | 0.420 |
|  | C | 0 | 0.1 | 0.1 | 0.8 | 0.005 | 0.001 | 0.098 | 0.076 | 0.824 | 0.399 |
|  | D | 0 | 0 | 0 | 1 | 0.008 | 0.001 | 0.004 | 0.021 | 0.974 | 0.006 |
| 2,000 | A | 0 | 0.03 | 0.02 | 0.95 | 0.005 | 0.000 | 0.028 | 0.005 | 0.967 | 0.417 |
|  | B | 0 | 0.1 | 0.02 | 0.88 | 0.006 | 0.000 | 0.098 | 0.007 | 0.895 | 0.429 |
|  | C | 0 | 0.1 | 0.1 | 0.8 | 0.006 | 0.000 | 0.098 | 0.001 | 0.901 | 0.450 |
|  | D | 0 | 0 | 0 | 1 | 0.010 | 0.000 | 0.000 | 0.000 | 1.000 | 0.001 |

**Table 3:**
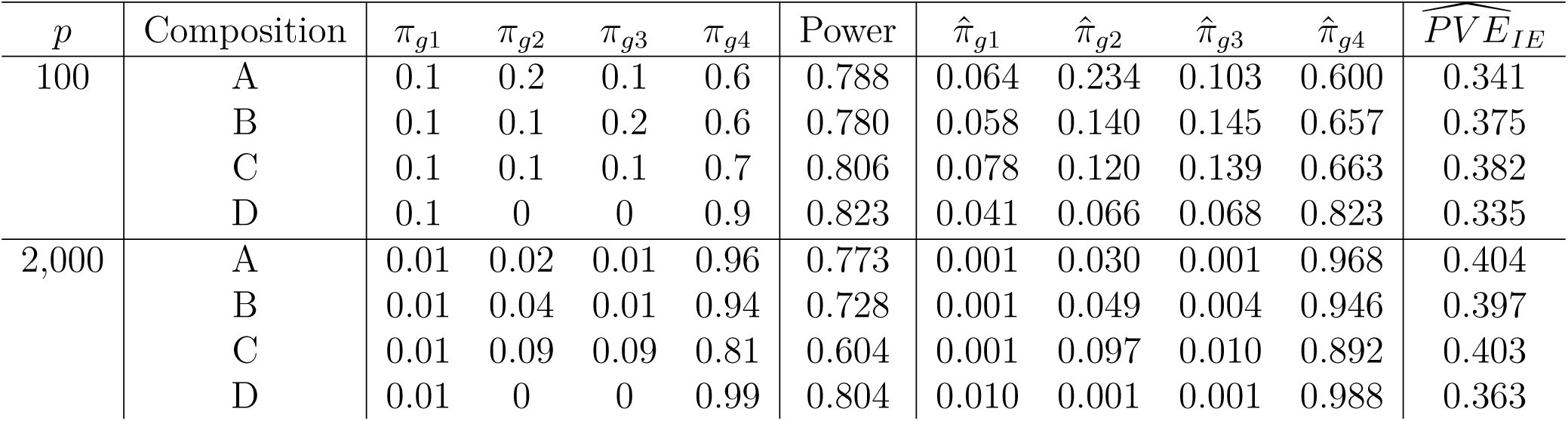
Empirical power of the proposed global test at level of 0.01 based on 1,000 simulations when *p* = 100/2, 000. We denote *π*_*g*1_, *π*_*g*2_, *π*_*g*3_ and *π*_*g*4_ to represent the proportion of mediators in Group 1, Group 2, Group 3 and Group 4 as defined in Table 1, and 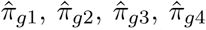 are the estimated proportions from our Bayesian method. The true *PVE_IE_* = 0.4, and 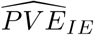 is the estimated value.

## 6 Data Analysis

We applied the proposed Bayesian method to investigate the mediation mechanism of DNAm in the pathway from adult socioeconomic status (SES) to glycated hemoglobin (HbA1c) in the Multi-Ethnic Study of Atherosclerosis (MESA) [31]. The exposure, adult SES, is indicated by adult educational attainment and is an important risk factor for cardiovascular diseases. The outcome, HbA1c, is a long-term measurement of average blood glucose levels and a critical variable for various diseases including T2D and CVD [32]. Thus, understanding how methylation at different CpG sites mediates the effects of adult SES on HbA1c can shed light on the molecular mechanisms of CVD. We provide our detailed processing steps for MESA data in the Supplementary Materials. Briefly, we selected 1,231 individuals with both adult SES and HbA1c measurements as well as DNA methylation profiles measured from purified monocytes. Due to computational reasons, we focused on a final set of 2,000 CpG sites that have the strongest marginal associations with adult SES for the following mediation analysis.

We applied both univariate mediation analysis and our Bayesian multivariate mediation analysis to analyze the selected 2,000 CpG sites. For the multivariate analysis, we consider

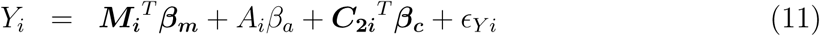

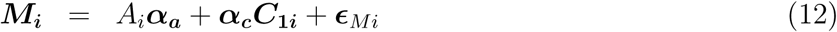

where *Y_i_* represnts HbA1c levels; *A_i_* represents adult SES values; and *M_i_* represnts methylation level for 2,000 CpG sites. In Equation (11), the model controls for age, gender and race/ethnicity, and in Equation (12), we adjust for age, gender, race/ethnicity and enrichment scores for 4 major blood cell types (neutrophils, B cells, T cells and natural killer cells). All the continuous variables are standardized to have zero mean and unit variance. The univariate analysis is applied in a similar fashion except that it is used to analyze one site at a time.

To compare the power between the univariate analysis and our Bayesian mediation analysis, we perform 100 permutations to obtain an empirical null distribution of PIP values following the procedure in Section 4.4, with which we obtain empirical estimates of FDRs for both univariate and Bayesian multivariate analysis. Consistent with simulations, our Bayesian multivariate mediation method is more powerful than the univariate mediation method. For example, at an FDR of 0.05, our Bayesian mediation method is able to identify 406 mediators while the univariate method is only able to identify 137. At an FDR of 0.10, our Bayesian mediation method can identify 612 mediators while the univariate mediation method can only identify 356.

We display PIP values for each of the 2,000 CpG sites from the Bayesian multivariate analysis in Figure 4. Two CpG sites were identified with strong evidence (PIP > 0.5) for mediating the adult SES effects on HbA1c. These two CpG sites are also among the top ten sites with the smallest *p*-values obtained from univariate mediation analysis. In addition, these two CpG sites are close to genes *CCDC54* and *CCND2*, both of which are known candidates associated with HbA1c. Specifically, the expression of *CCND2* has been shown to be associated with risk of T2D and the related glycemic traits of glucose, HbA1c, and insulin [33]. The gene *CCDC54* interacts with valproic acid and acrylamide, both of which are associated with diabetes and blood insulin [34, 35]. Therefore, strong evidence suggests that adult SES may act through these two genes to affect HbA1c.

**Figure 4:**
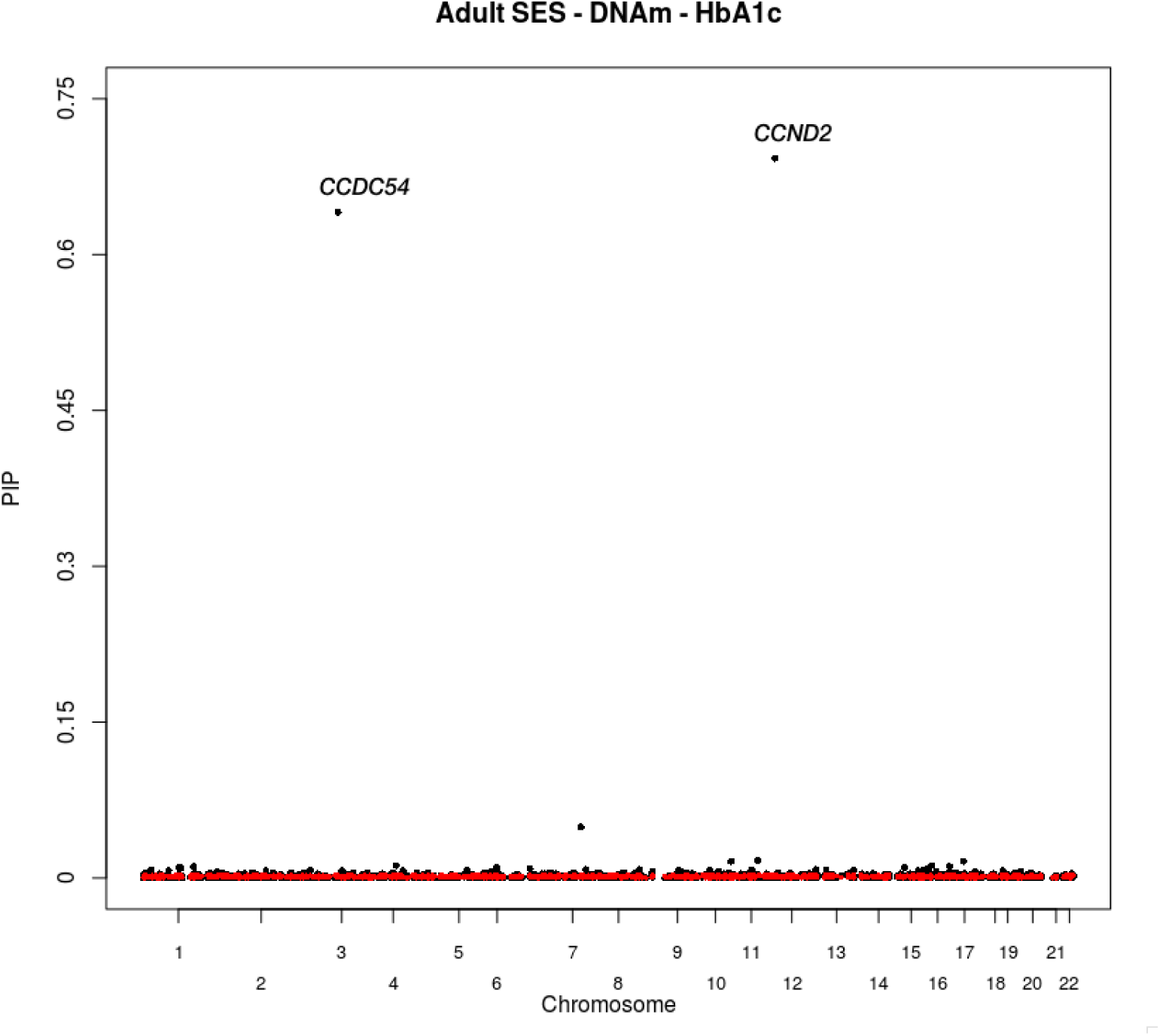
Consider the trio: Adult SES → DNAm → HbA1c. The black dots are the estimated posterior inclusion probability (PIP) for each CpG site from the Bayesian mediation method and the red dots are the estimated PIPs when we permute the outcome once and fit the Bayesian mediation method.

We also estimate the global mediation effects 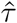 as 0.0084 and its 95% credible interval from the posterior samples as (0.0063, 0.0115). The *p*-value of the global test based on the empirical null distribution is 0.02. And the 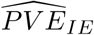is 0.096, indicating that approximately 10% of the outcome variance is indirectly explained by DNAm after controlling for covariates. In addition, we estimate the proportion of CpG sites in each of the four categories as defined in Section 4.3: 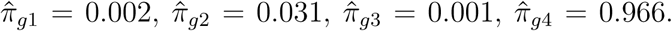 We find that a small proportion of DNAm has large effects on the HbA1c level, and a small proportion of DNAm is notably associated with adult SES. The results also suggest that adult SES acts through certain important DNAm sites to influence HbA1c.

## 7 Discussion

In this paper, we develop a Bayesian sparse linear mixed model for high-dimensional mediation analysis. The advantage of a Bayesian method is to propagate uncertainty for functions of parameters in a natural way instead of resorting to Delta methods or two-step approaches. Our method can simultaneously analyze a large number of unordered mediators without making any causally ordering assumptions. By imposing continuous shrinkage priors on the key regression coefficients for the mediation analysis, our method achieves up to 30% power gain in identifying true non-null mediators compared with univariate mediation method and approximately 10% power pain compared with multivariate method based on simulations. The Bayesian method also provides better interpretations of the way in which a mediator links or does not link exposure to outcome, and we construct tests for global indirect effects based on the structure of the composite null hypothesis. Our global test is slightly conservative under the null and yields decent power under the alternatives. Implementing our method to MESA, our Bayesian mediation method can detect more active mediators than univariate mediation method at fixed FDR levels. We also identified two genes, *CCDC54* and *CCND2*, with strong evidence for actively mediating the adult SES effects on HbA1c. Both of them are candidate genes associated with diabetes and blood insulin.

Recent literature proposes a convex penalty on the product term of indirect effect [36], which improves power of pathway selection and reduces estimation bias in the indirect effects. Under the Bayesian framework, direct shrinkage on the product term may be a more appropriate choice, as it takes into account the correlation between the two models in the mediation analysis and is more straightforward when the goal is to identify non-null mediators. Directly incorporating the correlation between the mediators will be another avenue to pursue. We leave that for future work.

